# Digital imaging to evaluate root system architectural changes associated with soil biotic factors

**DOI:** 10.1101/505321

**Authors:** Chakradhar Mattupalli, Anand Seethepalli, Larry M. York, Carolyn A. Young

## Abstract

Root system architecture (RSA) is critical for plant growth, which is influenced by several edaphic, environmental, genetic and biotic factors including beneficial and pathogenic microbes. Studying root architecture and the dynamic changes that occur during a plant’s lifespan, especially for perennial crops growing over multiple growing seasons, is still a challenge because of the nature of their growing environment in soil. We describe the utility of an imaging platform called RhizoVision Crown to study RSA of alfalfa, a perennial forage crop affected by Phymatotrichopsis Root Rot (PRR) disease. *Phymatotrichopsis omnivora* is the causal agent of PRR disease that reduces alfalfa stand longevity. During the lifetime of the stand, PRR disease rings enlarge and the field can be categorized into three zones based upon plant status: asymptomatic, disease front and survivor. To study root architectural changes associated with PRR, a four-year old 25.6-hectare alfalfa stand infested with PRR was selected at the Red River Farm, Burneyville, OK during October 2017. Line transect sampling was conducted from four actively growing PRR disease rings. At each disease ring, six line transects were positioned spanning 15 m on either side of the disease front with one alfalfa root sampled at every 3 m interval. Each alfalfa root was imaged with the RhizoVision Crown platform using a backlight and a high-resolution monochrome CMOS camera enabling preservation of the natural root architectural integrity. The platform’s image analysis software, RhizoVision Analyzer, automatically segmented images, skeletonized, and extracted a suite of features. Data indicated that the survivor plants compensated for damage or loss to the taproot through the development of more lateral and crown roots, and that a suite of multivariate features could be used to automatically classify roots as from survivor or asymptomatic zones. Root growth is a dynamic process adapting to ever changing interactions among various phytobiome components, by utilizing a low-cost, efficient and high-throughput Rhizo-Vision Crown platform we showed quantification of these changes occurring in a mature perennial forage crop.

## INTRODUCTION

Roots function at the soil interface and encounter complex phytobiome components in a dynamic environment. Root architecture plays an important role in exploiting unevenly distributed soil resources such as water and nutrients, which in turn determines plant productivity (Lynch 1995; Paez-Garcia et al. 2015). For example, maize genotypes with fewer crown roots growing in low nitrogen soils had greater rooting depth, captured more nitrogen from deep soil layers and had greater relative yield compared to genotypes with more crown roots (Saengwilai et al. 2014). Similarly, root crown properties such as root number, angles, and complexity have been correlated to plant performance and rooting depth in the field, because the root crown serves as the backbone of the root system (Saengwilai et al. 2014; Trachsel et al. 2013; York and Lynch 2015). By nature of being in the soil, evaluating roots undergoing biotic or abiotic stresses can be inherently difficult, especially when evaluating field grown plants that exhibit robust three-dimensional root structures.

Several approaches have been used in the past to study various root morphological and architectural parameters. These included tracings using modified line-transect methods to estimate root lengths, taking soil cores to determine root length densities, or water-displacement method for assessing root volume (Hancock 1991; Harrington et al. 1994; Larkin et al. 1995). A major limitation with these methods is their inability to provide any information about root system architecture. Acquiring digital root images followed by their analyses using software programs such as ROOTEDGE or WinRHIZO™ has been used to assess root growth characteristics such as total length, surface area, volume, number of links, average radius and other topological features (Arias et al. 2013; Himmelbauer et al. 2004; Ma et al. 2014; Ortiz-Ribbing and Eastburn 2004). However, these systems are often not suitable for studying three-dimensional, complex root samples from perennial field grown plants.

In order to overcome the challenges presented by mature root systems, root crown phenotyping (York 2018), also called shovelomics, was developed. Shovelomics involves excavating the top portion of the root system, here defined as the root crown, washing to remove soil, and then quantifying root system architectural parameters. Shovelomics originally used visual scoring to study basic architectural traits of mature maize root crowns in the field (Trachsel et al. 2011). This approach was further extended by an imaging protocol and algorithmic approach, where the excised roots were positioned on a black background with diffuse reflectance properties and photographs were taken using a digital camera mounted on a tripod (Bucksch et al. 2014). Further refinement of this digital root imaging process was achieved through the development of RhizoVision Crown, a phenotyping platform that integrates open hardware and software to streamline measurements of root crowns excavated from the field. The hardware component of the RhizoVision Crown is an enclosed imaging box with an LED backlight on one side, and a monochrome CMOS camera on the other, which allows the silhouette of the root crown to be captured. Use of the silhouette facilitates downstream image analysis by making identification of the root object easily achieved using simple threadholding. A suite of traits are computed including root numbers, diameters, root crown size, angles, and total length, many of which can be influenced by biotic and abiotic stresses.

Considerable research has been made to evaluate how biotic stresses can induce changes in root morphology and architecture. For example, infection of greenhouse grown alfalfa seedling roots by *Pythium ultimum* and *Pythium irregulare* resulted in smaller root system size and complexity with a lower degree of branching compared with non-infected plants (Larkin et al. 1995). Reduction in total root length, fresh and dry weight of roots was observed when crops such as bermudagrass and onion were infected with plant-parasitic nematodes (Luc et al. 2006; Pang et al. 2009). Similarly, cotton seedlings infected with either a fungus (*Thielaviopsis basicola*) or a root-knot nematode (*Meloidogyne incognita*) led to a reduction in root fresh weight as well as root morphological (total root length, root surface area, total number of root links) and topological (magnitude, altitude and exterior path length) parameters (Ma et al. (2014). However, many of these studies have been based on short-term experiments with younger plants that have more malleable root systems that facilitate easier evaluation.

We evaluated the utility of the RhizoVision Crown imaging platform for studying the effects of a biotic factor, *Phymatotrichopsis omnivora* causal agent of Phymatotrichopsis Root Rot (PRR) disease on alfalfa. Alfalfa (*Medicago sativa* L.) is the leading perennial forage legume crop in the USA. Profitable alfalfa production in western United States and Mexico can be greatly affected by PRR disease (also known as Phymatotrichum root rot, cotton root rot, Texas root rot and Ozonium root rot). PRR affected alfalfa plants exhibit roots with discolored to dark brown vascular tissues, and necrotic lesions with dead cortical tissues that slough off readily. Above ground, diseased plants observed during mid to late summer wilt rapidly followed by death with leaves firmly attached to the stem (Fig. 1A). At a landscape level, PRR spreads in a centrifugal fashion forming numerous circular to irregular shaped disease foci (Fig. 1B) (Mattupalli et al. 2017; Streets and Bloss 1973; Uppalapati et al. 2010; Young et al. 2015). During multiple growing seasons, these disease foci expand and some alfalfa plants in the center recover and re-establish, but a comprehensive knowledge of the root architectural changes in response to PRR is lacking (King 1923; Mattupalli et al. 2018). Hence, the objective of this study was to test the hypothesis that alfalfa plants surviving from PRR disease stress would have root systems acclimated to damage caused by the disease. This hypothesis was tested using the RhizoVision Crown phenotyping platform and comparisons were made between root crowns of asymptomatic and PRR disease survivor alfalfa plants sampled from a PRR-infested four-year old alfalfa stand.

**Figure 1:**
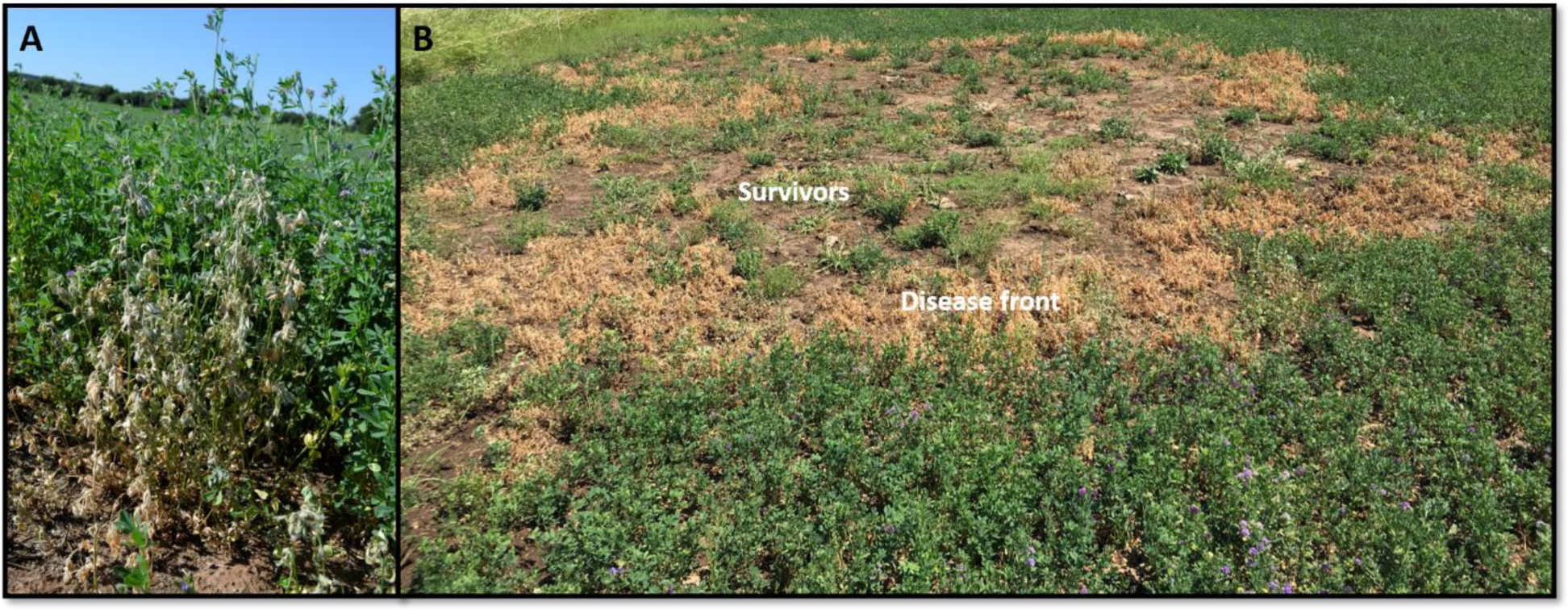
Observation of Phymatotrichopsis root rot (PRR) symptoms in the field. A) Alfalfa showing PRR disease symptoms. Diseased plants exhibit rapid wilting with leaves attached to the stem. B) Example of a PRR disease ring. Circular to irregularly shaped PRR disease foci showing dead alfalfa plants at the disease front and survivors in the center.

## MATERIALS AND METHODS

### Field site history

The study was conducted on a PRR-infested 25.6 ha semi-circular alfalfa commercial hay production field under center-pivot irrigation located at the Noble Research Institute’s Red River Farm, Burneyville, Oklahoma (Fig. 2A; 33°52’36.0”N, 97°15’43.0”W) that was previously a native pecan orchard. The soil type in this field was categorized as gaddy loam to Yahola fine sandy loam. The orchard was removed and planted with soybean and small grains during 2005-2013 followed by America’s Alfalfa Alfagraze 600 RR in the autumn of 2013. The alfalfa stand was maintained following commercial hay production agronomic management practices followed in the southern Oklahoma region.

**Figure 2:**
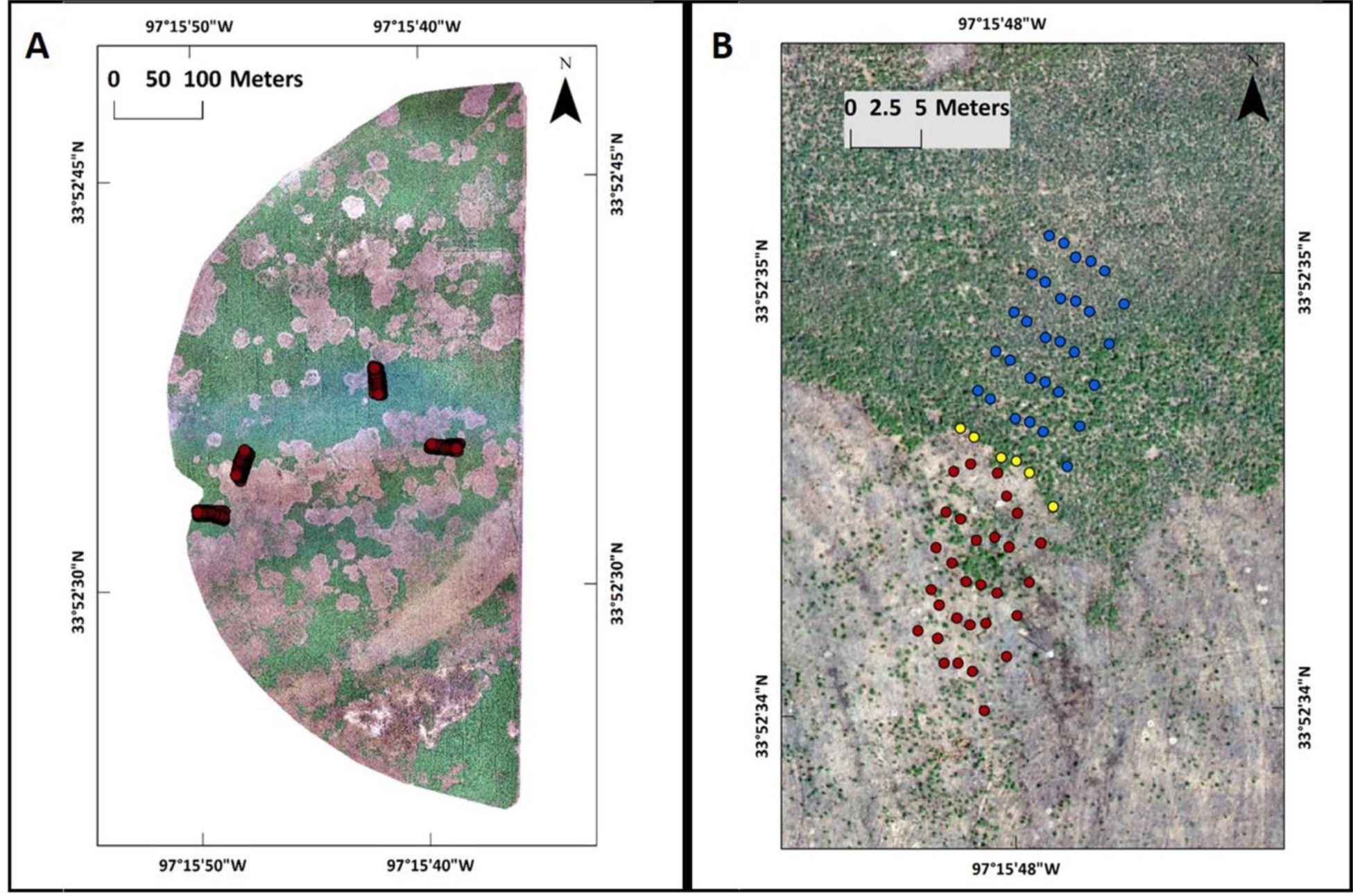
Alfalfa field study site. A) Alfalfa field infested with Phymatotrichopsis root rot (PRR) disease located near Burneyville, OK. Red dots represent four sampling PRR disease rings. B) A representative PRR disease site showing line transect sampling procedure. Red, yellow and blue dots indicate the locations where alfalfa root crowns were sampled from survivor, disease front and asymptomatic zones respectively.

### Sampling strategy

Sampling of alfalfa root crowns was performed during October 2017, a time period coinciding with onset of alfalfa fall dormancy. As the disease progresses through the field, a typical PRR disease ring can be classified into three zones based on visual observations: a) a strip of dead plants at the disease front zone, which differentiates survivor and asymptomatic zones b) a survivor zone extending inwards from the disease front zone, where most plants have succumbed to PRR disease with some plants re-establishing as survivors c) an asymptomatic zone extending outwards from the disease front zone, with alfalfa plants showing no visible disease symptoms. At each of the four PRR-disease rings, sampling was performed along six line transects that would encompass disease progression over at least two growing seasons (Fig. 2B). Along each line transect one plant was sampled every 3 m interval spanning both asymptomatic and survivor zones with the disease front as the midpoint to a total of 30 m. Root crowns in the disease front zone were sampled from the first unaffected plant based on visual inspection. Alfalfa plants were dug out of the soil using a shovel focusing on obtaining roots from the top 20-25 cm of soil, which represented the crown and provided information on the integrity of the tap root. There were instances where no alfalfa plant was present at the exact sampling distances (3, 6, 9, 12, 15 m) in the survivor zone. In such cases, the plant nearest to the intended sampling distance was sampled. The root crowns were separated from the above-ground foliage and soil was brushed off the roots and imaged in the laboratory using the RhizoVision Crown platform. Roots were also scored visually for the presence of lesions and necrotic or loss of taproots due to PRR disease.

### RhizoVision Crown hardware platform

The RhizoVision Crown hardware platform (Fig. 3) consists of T-slotted aluminum profiles (80/20 Inc., Columbia City, IN) that were assembled to generate a box measuring 65.5 cm x 65.5 cm x 91.4 cm. Foamed black PVC panels (TAP Plastics, Stockton, CA), were inserted between profiles to isolate the interior from outside light. On one end, a 61 cm x 61 cm LED edge lit flat panel light (Anten, 40 watts, 6000K light color) was affixed using epoxy. Across and centered, a CMOS sensor monochrome camera (Basler acA3800-14um, Graftek Imaging, Inc., Austin, TX) was mounted. The camera requires a single USB 3 cable that transfers data and supplies power. The camera was attached to a laptop computer with a USB barcode scanner (Tautronics, Fremont, CA) installed. The software components include RhizoVision Imager that controls the camera, and RhizoVision Analyzer that extracts measurements from the images. Imager software was used to set the camera properties to an exposure time of 10,000 µs and a gamma of 3.9 to increase contrast. Files were named manually in the Imager software before acquiring the image and storing on the computer.

**Figure 3:**
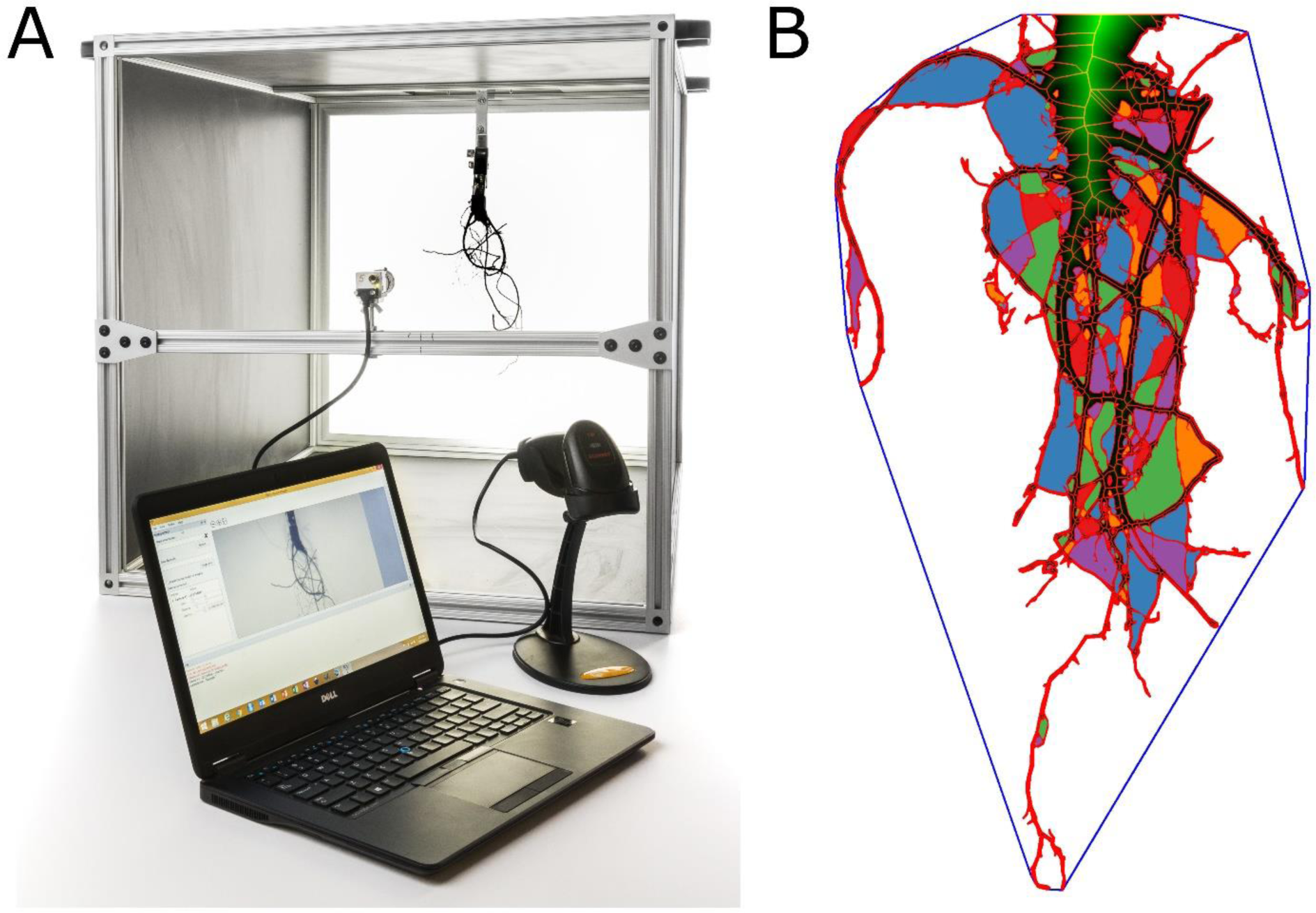
RhizoVision Crown platform. A) Components of the RhizoVision Crown platform. B) A representative segmented image of a root crown taken by the RhizoVision Crown platform including the distance map (green heat map) and medial axis skeleton (central red lines), root perimeter (red outline), the holes within the root crown (multi-colored to aid separation), and the convex hull (blue line surrounding whole root crown).

### Extraction of Features

Image features are measurements extracted using image analysis. RhizoVision Analyzer software was utilized in this study. Analyzer works in batch mode to process a folder of files to output a data CSV file and has options to output feature images that include the derived metrics overlaid on the segmented image. Segmentation is by simple thresholding of the greyscale values for each pixel due to the optimization of the imaging method. Following segmentation, the edges are smoothed to remove small irregularities that falsely contribute to root length. This smoothed image is used for calculating the surface area, the volume, and the perimeter, number of holes (disconnected components of inverted image), average hole size, the convex hull, and solidity (Table 1). Next, the smoothed image is skeletonized using a distance transform followed by identifying the medial axes. From this skeleton, total root length, diameters, and angles are calculated. Based on the diameter of the medial axis pixels in the skeletonized images, the roots in each image were categorized into fine (< 1.7 mm), medium (1.7 – 3.41 mm) or coarse root (> 3.41 mm). The orientation in degrees from horizontal of all skeleton root pixels was determined based on a 20-pixel window around each respective pixel and categorized into steep (> 60 °), medium (60-30 °), or shallow (< 30 °).

**Table 1:** Description of various features extracted from digital images in the study.

| Features extracted<sup>a</sup> | Description |
| --- | --- |
| Median and maximum number of roots | The number of roots was counted by performing horizontal line scans from left to right in each row through the segmented image. In each of the line scan, we check if there is a pixel value transition from the previous pixel value to the current pixel value on its right side. |
| Number of root tips | Tips were defined as the end-point pixels of the skeletonized image. |
| Total root length | Computed by calculating Euclidean distances along the skeletonized image. |
| Depth, maximum width and width-to-depth ratio | Depth and maximum width of the root in the segmented image. The ratio of maximum width to depth of the image was noted as width-to-depth ratio. |
| Network area, convex area and solidity | Network area is based on the total number of pixels in the segmented image. The convex hull of a geometric shape is minimal sized convex polygon that can contain the shape. The ratio of network area and the convex area was noted as the solidity. |
| Perimeter | Perimeter is based count of total number of pixels in the perimeter image (outlining the roots). |
| Average, Median and maximum diameter | For each pixel on the skeletonized image, the distance to the nearest non-root pixel was computed and noted as the radius at that pixel. The list of radii from all the medial axis pixels was obtained to determine the average and maximum diameter. |
| Volume and surface area | Using the radii determined earlier, the sum of all cross-sectional areas across all the medial axis pixels were noted as volume and the sum of the perimeter across all the medial axis pixels were noted as surface area. |
| Lower root area | The lower root area was the area of the segmented image pixels that were located below the location of the medial axis pixel that had the maximum radius. |
| Holes | Holes were the disconnected background components and indicative of root branching and complexity. They can be counted by inverting the segmented image. The average hole size (area) is also calculated. |
| Fine diameter frequency,<br>Medium diameter frequency,<br>Coarse diameter frequency | From the skeletal image, the medial axis pixels were grouped into fine or coarse roots based on the radius values at the pixels. |
| Root angle orientation frequencies | Calculated by observing the slope of 20-pixel window around each pixel of the skeletonized image and binning as steep, medium, or shallow. |
| Computational time | The time taken to extract traits for every plant root image. |

<sup>a</sup> Figure 3B shows a representative root crown image from which various root features were extracted

### PCR assay for *P. omnivora* detection

*P. omnivora* presence in the root was assessed by obtaining two sub-samples from each sampled alfalfa root. The tissue was finely chopped using a scalpel blade and lyophilized prior to DNA extraction. DNA was extracted using MagAttract 96 DNA Plant Core Kit (Qiagen Inc., Valencia, CA) following manufacturer’s protocol. Detection of *P. omnivora* from these samples was carried out in an end-point PCR assay with PO2F/PO2R primers that were previously developed by Arif et al. (2014). Each reaction consisted of 3 µl of extracted DNA, 0.5 µl of each 10 µM primer, 5 µl of 5X Green GoTaq^®^ reaction buffer (Promega Corporation, Madison, WI), 0.5 µl of 10 mM dNTP mix (Promega Corporation, Madison, WI), and 0.2 µl of GoTaq^®^ DNA Polymerase (Promega Corporation, Madison, WI) in 25 µl of final volume. All reactions were run on Applied Biosystem 2720 Thermal Cycler using the following protocol: 95°C for 2 min followed by 35 cycles of 95°C denaturation for 15 s, annealing at 60°C for 30 s, extension at 72°C for 45 s, and a final elongation at 72°C for 7 min. *P. omnivora* mycelial DNA (1 ng/ µl) and sterile water were used as positive and negative controls, respectively. PCR products (10 µl) were then subjected to electrophoresis in 1.5% agarose gel at 80 Volts for 90 min, stained with ethidium bromide and visualized in UVP GelDoc-It imaging system. An alfalfa root was considered positive for *P. omnivora* if the pathogen was detected in one or both sub-samples.

### Data analysis

Data was analyzed using R (3.5.1). Plots were created using ggplot2 package 3.0.0 (Wickham 2016). Means and standard errors were calculated for all features using the dplyr package 0.7.6 (Wickham et al. 2018). ANOVA was conducted using the ‘aov’ function to test for the effect of the zone ID and distance for every trait. Distance was found to have no significant effects, so for subsequent analysis only zones were used as explanatory factors. The site was considered as a block for the error term. ANOVAs were conducted using all three zones and with the three combinations of pairwise zone combinations, which allowed us to look at the effect of dropping zones. The disease front zone was found to have more significant differences with the survivors than asymptomatic plants, and therefore was removed for further analysis given that it has fewer samples. Pairwise correlation analysis was conducted for all pairwise combinations of image features using the ‘corr’ function. Principal component analysis was conducted with missing values omitted using the ‘prcomp’ function using centered and scaled data. Linear discriminant analysis was conducted using the ‘lda’ function from the MASS package to predict the survivor or asymptomatic classes using data that was standardized for each measurement such that the mean was zero and the within-group standard deviation was one in order to interpret loadings (Venables and Ripley 2002).

## RESULTS

Sampled root crowns were visually assessed for taproot status and the results are presented in Figure 4. All root crowns from the asymptomatic zone had a taproot with no evidence of lesions, whereas 54-79 % of root crowns sampled from survivor zone exhibited taproots that were either missing completely or sloughed off partially. A lower percentage of root crowns (8-17 %) from survivor and disease front zones had evidence of a necrotic taproot. To detect *P. omnivora,* alfalfa root crowns sampled from the study site were subjected to an end point PCR using molecular markers specific for the pathogen. The results were expressed as a percentage of root crowns positive for *P. omnivora* presence (Fig. 5). Data indicated that a higher percentage (75%) of root crowns sampled from the disease front and those within 6 m of the survivor zone were positive for the presence of the pathogen. The pathogen detection percentage decreased as sampling distance increased from the disease front into the survivor zone. Even though plants in the asymptomatic zone did not show any visual symptoms of PRR disease, we were still able to detect the pathogen in some root samples, even out as far as 15 m from the disease front.

**Figure 4:**
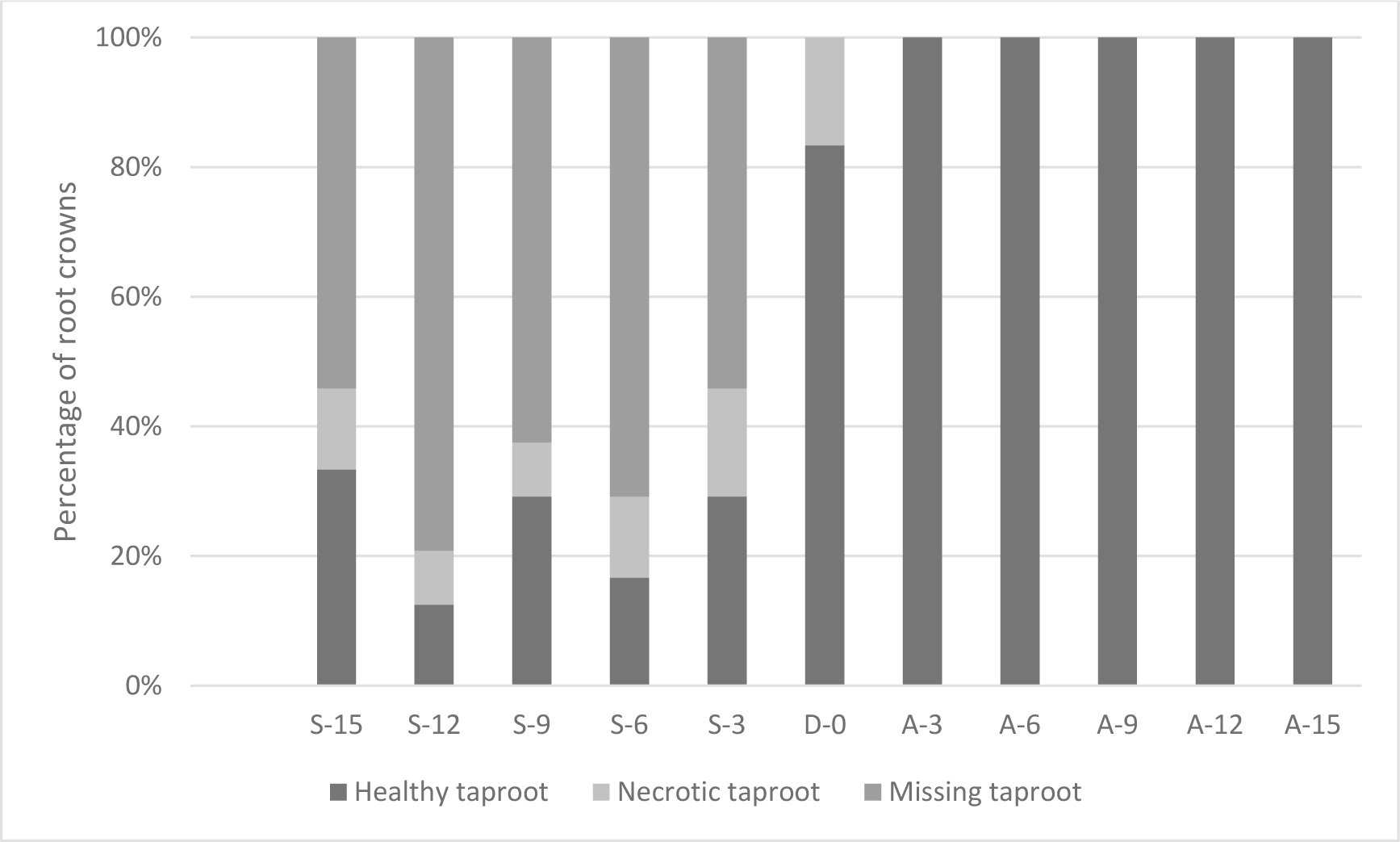
Taproot status of alfalfa root crowns sampled from a Phymatotrichopsis root rot infested field near Burneyville, OK. S, D and A represent samples from symptomatic, disease front and asymptomatic zones, respectively. Roots were visually examined for their integrity, based on the presence of healthy taproot, necrotic taproot or missing taproot. Letters followed by numbers show the distance (in meters) from disease front.

**Figure 5:**
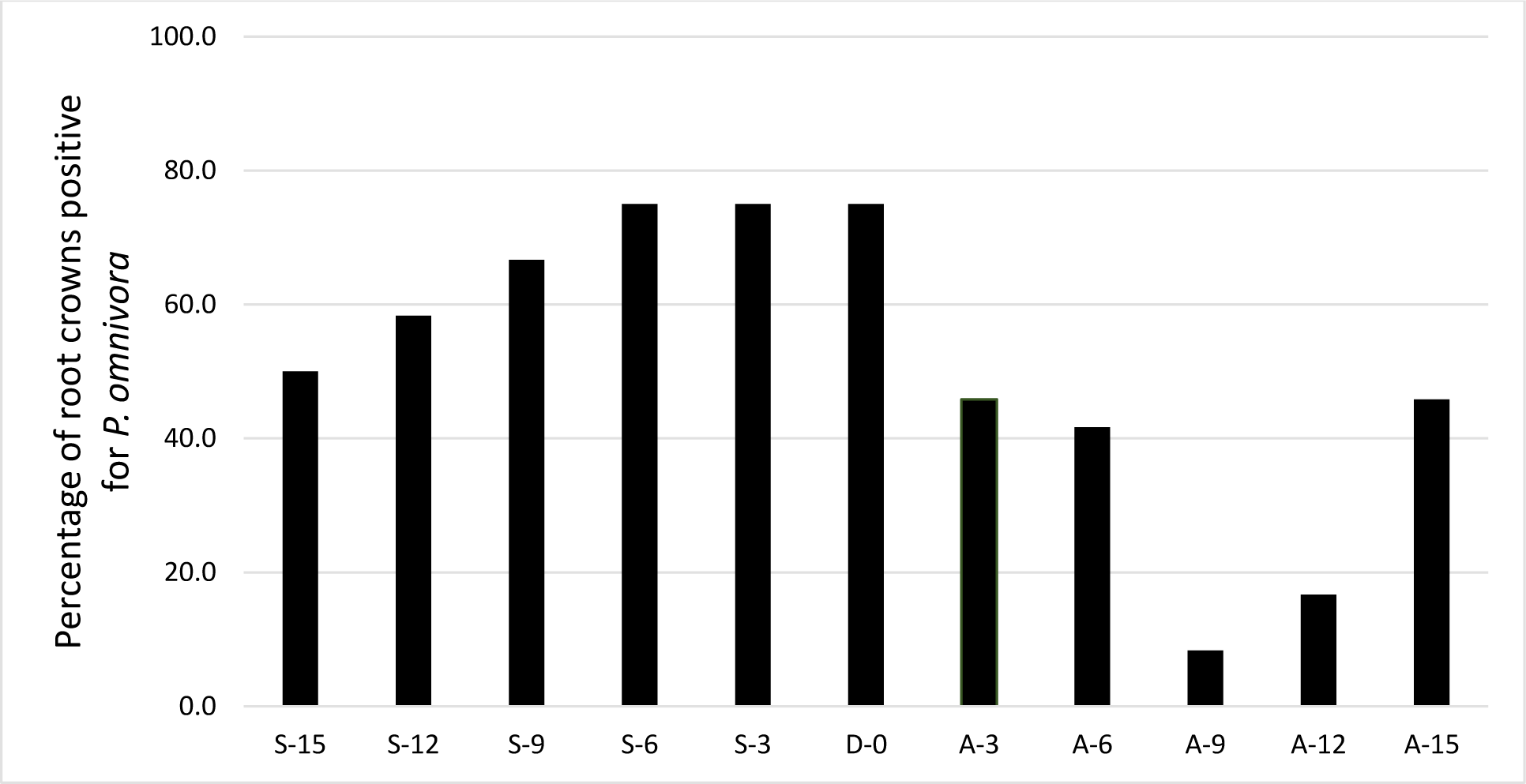
PCR detection of *P. omnivora* from alfalfa root crowns sampled from a Phymatotrichopsis root rot infested field near Burneyville, OK. S, D and A represent samples from symptomatic, disease front and asymptomatic zones, respectively. Letters followed by numbers show the distance (in meters) from disease front.

RhizoVision Analyzer software was developed to extract an array of features from root crown images obtained using the RhizoVision Crown imaging platform. The line transect sampling strategy resulted in sampling of a small number of root crowns from the disease front zone (n=24) compared to sample size of the asymptomatic or survivor zones (n=120 from each zone). During preliminary ANOVA analysis, root crowns of asymptomatic and disease front zones differed significantly for only one of the 27 features assessed, demonstrating that the disease front root crowns were more or less indistinguishable from asymptomatic zone root crowns (data not shown). Hence, further analyses were focused only on root crowns from the survivor and asymptomatic zones.

The means and standard errors of the features extracted from alfalfa root crown images that were sampled from survivor and asymptomatic zones are shown in Figure 6. Root crowns sampled from the survivor zone (17.3 ± 0.68) had greater numbers of roots compared to samples from the asymptomatic zone (15.2 ± 0.61). Likewise root crowns from the survivor zone had more total root length (2531.3 ± 142.05 mm), perimeter (2991.3 ± 157.86 mm), root depth (161.4 ± 5.41 mm), root width (129.7 ± 4.78 mm), and root tips (406.7 ± 20.38) than those from asymptomatic zone (2043.5 ± 110.33 mm; 2399.3 ± 117.57 mm; 143.3 ± 4.37 mm; 117.2 ± 3.65 mm; 301.9 ± 16.6, respectively). However, there was no significant difference in root width to depth ratio between root crowns from asymptomatic and survivor zones.

**Figure 6:**
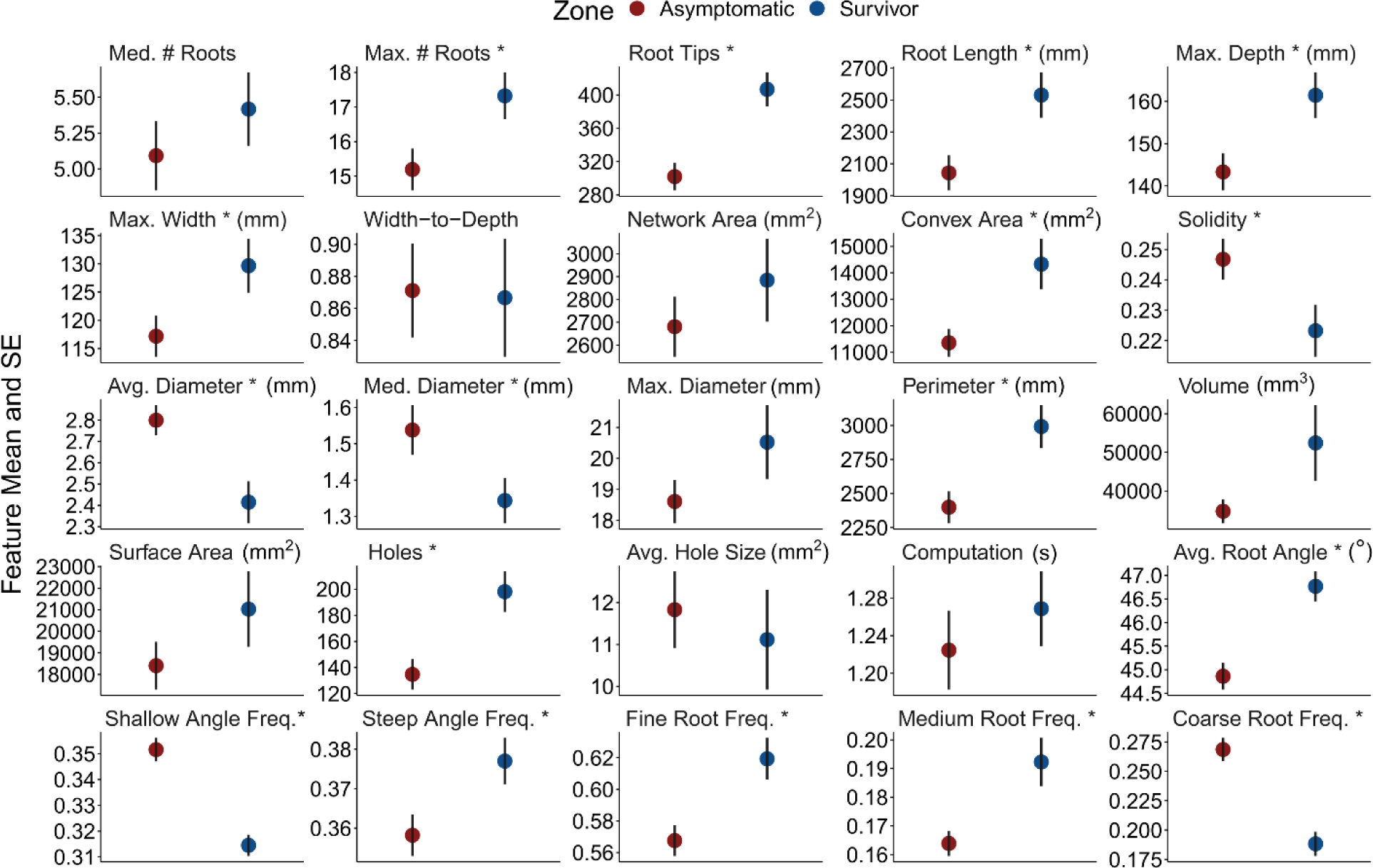
Summary of means and standard errors of various features extracted from alfalfa root crown images that were sampled from survivor and asymptomatic zones of four Phymatotrichopsis root rot disease rings. Features that showed significant differences between the two zones are noted with an asterisk.

Network area was not significantly different between root crowns of asymptomatic and survivor zones (Figure 6). Convex area was significantly greater in root crowns from survivor zone (14329 ± 960.93 mm^2^) than those from asymptomatic zone (11353 ± 530.41 mm^2^). Interestingly the solidity feature, which was calculated as ratio of network and convex areas, was significantly greater for roots from asymptomatic zone (0.25 ± 0.01) than the root crowns sampled from survivor zone (0.22 ± 0.01).

There was no difference between maximum diameter of root crowns from survivor and asymptomatic zones, but the average and median root diameter of asymptomatic zone root crowns was significantly different (2.8 ± 0.07 mm; 1.54 ± 0.07 mm) from survivor zone roots (2.4 ± 0.1 mm; 1.34 ± 0.06 mm). Total root volume, lower root area and surface area were not different between the two zone root crowns. Roots from the asymptomatic zone had significantly higher frequency of coarse roots (0.27 ± 0.01), while higher frequency of fine (0.62 ± 0.01) and medium roots (0.19 ± 0.01) roots were observed with roots from the survivor zone. The average root angle was significantly steeper in the survivor zone than asymptomatic (46.8 ± 0.32, 44.9 ± 0.28, respectively). Root crowns sampled from the asymptomatic zone had significantly higher frequency of shallow angled roots (0.35 ± 0.005) compared to the survivor zone (0.31 ± 0.004). On the contrary, survivor zone root crowns had higher medium angle (0.31 ± 0.004) and steep angle (0.38 ± 0.006) frequency of roots than asymptomatic zone root crowns. The RhizoVision Analyzer software also calculated the number of disconnected background components in the image, described as holes, which is an indication of root structure complexity. Data suggested that root crowns from survivor zone had significantly more holes (198.1 ± 15.49) than the roots from asymptomatic zone (134.7 ± 11.63), but the average hole size did not differ. Finally, computation time required to extract all the features in the study was not different for the images obtained from either asymptomatic (1.22 ± 0.04 s) or survivor (1.27 ± 0.04 s) zones.

Pearson’s correlation coefficient was applied to find relationships between various features extracted from alfalfa root crown images and the results are shown in the form of a heatmap (Fig. 7A). A strong positive correlation was seen between total root length and the number of holes indicating root crowns with more complex roots systems have longer total root length. As expected, a negative correlation was observed between solidity and convex area.

**Figure 7:**
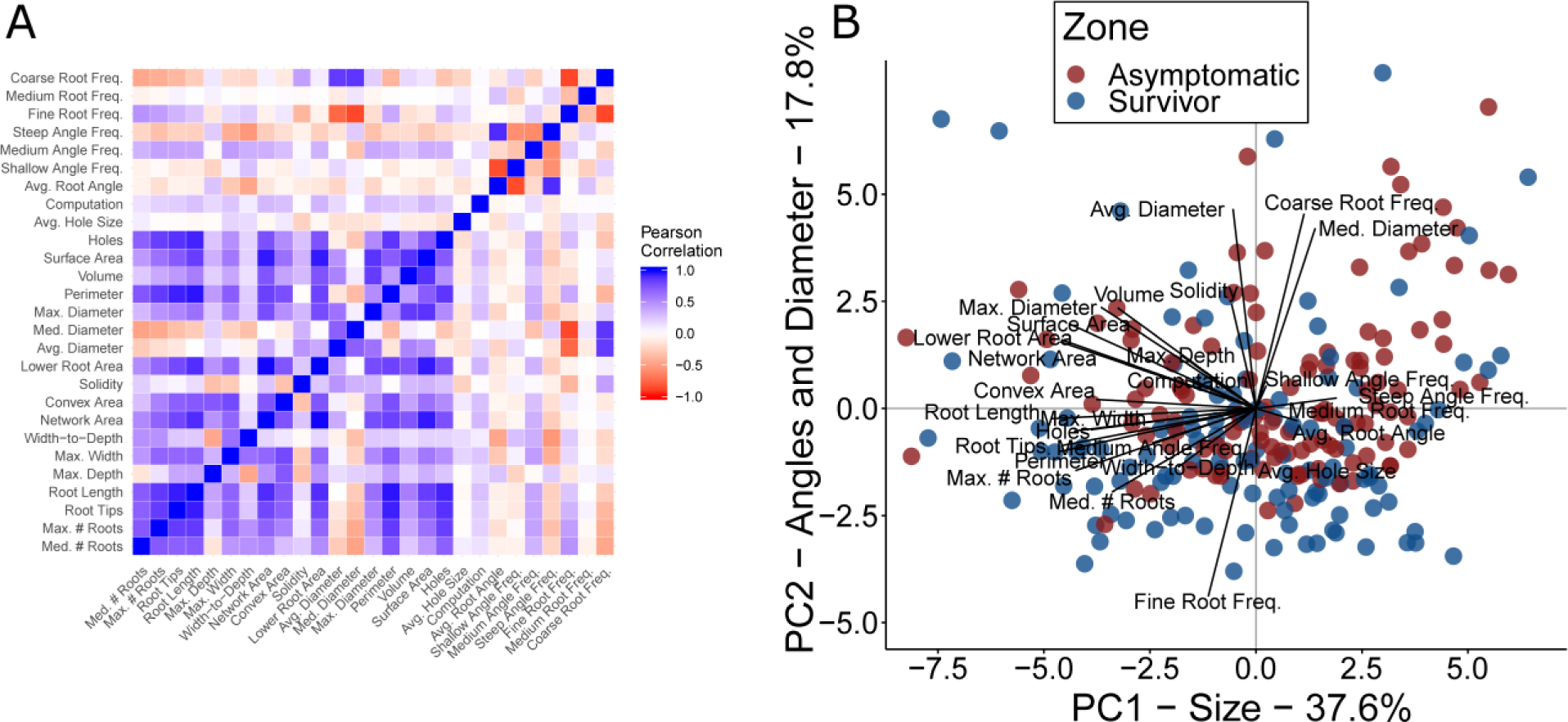
Relationship between various root features extracted using RhizoVision Analyzer software. Pearson’s correlation heatmap (A) and Principal component analysis (B) of root crown features.

In order to further investigate the correlation structure of these data, principal component analysis was used (Fig. 7B). The first and second components explained 37.6% and 17.8% of the multivariate variation of the root crowns. The first principal component was loaded most strongly by total root length, network area, number of root tips, holes, and numbers of roots, which are indicators of root system size. The second component was loaded most strongly by diameters and solidity, which can be viewed as indicators of root system exploration efficiency. Root crowns of asymptomatic and survivor zones separated more strongly along the second component, but overall separation was not substantial.

Linear discriminant analysis used the multivariate image feature data to maximize the separation of the survivor and asymptomatic zone classes (Fig. 8). Given two classes, one linear discriminant function was returned that provided an overall classification accuracy as either survivor or asymptomatic for individual root crowns of 75.6%. Because the data was standardized, the loadings of individual features in the discriminant function can be interpreted as the relative influence of any given trait on this classification ability. Total root length had the greatest absolute loading with a value of −8.5, other notable influencing features being surface area (4.0), perimeter (3.5), number of holes (2.3), and number of root tips (1.8), median diameter (1.1), with other features having loadings of one or less. Negative values indicate that the feature contributed to the asymptomatic class, while positive values indicate the feature contributed to the survivor class.

**Figure 8:**
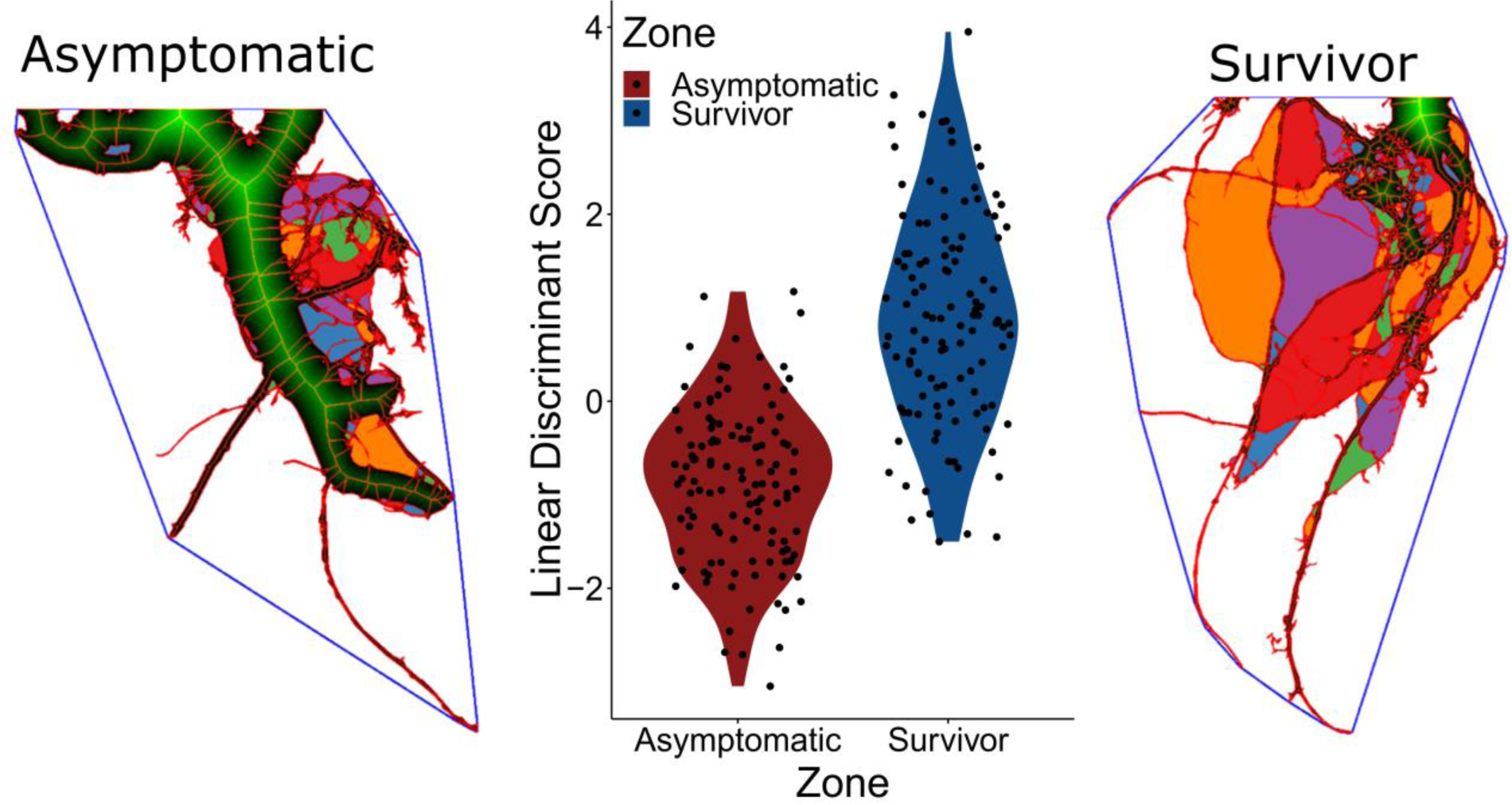
Linear discriminant analysis indicating differences between multivariate features extracted from alfalfa root crown images obtained from asymptomatic and survivor zones of four Phymatotrichopsis root rot disease rings. Representative images highlighting these differences are shown with extracted features overlaid on the original segmented root crown for asymptomatic (left) and survivor (right) plants.

## DISCUSSION

Several root phenotypes such as crown roots, differential production of hypocotyl-borne roots and root cortical aerenchyma have been identified in beans and maize that enhance greater acquisition of water and nutrients like nitrogen and phosphorus from soil while reducing the metabolic costs involved in soil exploration (Lynch 2015). However, such studies focused on annual crops and abiotic components of the phytobiome. Perennial crops like alfalfa also has strategies such as a taproot that can penetrate over six meters deep into soil enabling the plant to acquire water and nutrients from deeper soil layers, which makes it as one of the most drought-tolerant crops (Undersander et al. 2011). Yet, several root rot pathogens such as *Phytophthora medicaginis* and *Phoma sclerotioides* threaten the host reliance on the single taproot for long term survivability (Samac et al. 2015).

This study focused on evaluating roots from four-year-old alfalfa plants grown in a stand that was infested with the causal agent of PRR. PRR disease symptoms were initially noticed at the end of first growing year, but over time the stand became riddled with numerous disease foci, each year growing larger in size (Mattupalli et al. 2018). One interesting observation was the presence of surviving plants within the disease foci. In order to comprehensively document alfalfa root system architectural changes to PRR, we utilized the RhizoVision Crown platform and the RhizoVision Analyzer software. This approach demonstrated the ability to take digital images from mature root crowns and extract many root features enabling easy quantification of root system architecture.

Previous root studies analyzed images of young seedling roots or annual plant roots that were obtained by scanning in a flatbed scanner followed by using algorithms such as winRHIZO or ROOTEDGE (Arias et al. 2013; Himmelbauer et al. 2004). One of the key differences compared to using a flatbed scanner is that the RhizoVision Crown platform can preserve the integrity of mature root crowns which are more complex and often three dimensional. Scanning on a flatbed scanner would be problematic due to the inability of a mature root crown to lay flat on a planar surface. However, the RhizoVision Crown platform with an LED backlight and a monochrome CMOS camera has made acquiring images of four-year old alfalfa root crowns quick, reproducible, and ready for digital image analysis. Although the images were taken in a laboratory setting, RhizoVision Crown platform is portable thereby extending a researcher’s ability to acquire data directly in the field. While RhizoVision Crown optimizes imaging and analysis of root crowns, one of the limitations is that this platform lacks color information due to the backlit method.

Since *P. omnivora* was difficult to isolate from mature roots, a PCR-based approach was used to detect the pathogen in alfalfa root crown samples (Arif et al. 2014; Arif et al. 2013). *P. omnivora* was detected from a higher percentage of roots sampled from the survivor zone. PRR disease symptoms were not observed on plants from the asymptomatic zone, but the pathogen was detected from a limited number of root crowns sampled in this zone. This aligns with previous studies where *P. omnivora* was observed on 60-100% of the asymptomatic apple trees that were spaced 3-m^2^ apart from the symptomatic trees (Watson et al. 2000). Interestingly, in our study, an increase in the rate of detection was observed from samples obtained 9 m further away from the disease front into the asymptomatic zone. We hypothesize that this occurrence might be due to the proximity of plants at this distance to other nearby PRR disease rings as it was difficult to identify isolated disease rings.

From a temporal perspective, PRR manifests as diseased areas that expand in a circular to irregular shaped fashion, causing 31.4% reduction in alfalfa stand over two growing seasons (Mattupalli et al. 2018). Our sampling strategy was specifically developed to enable us to screen alfalfa plants that would have succumbed to the disease across different years. As the disease progresses, some alfalfa plants appear to cope with the damage or loss of the taproot and were able to reestablish as survivors. However, we don’t know if a plant has simply escaped disease until a plant crown is dug up and evaluated. In this study, several root system architectural differences were observed with root crowns sampled from the survivor zone compared to the asymptomatic zone.

A significant increase in the total root length, number of root tips, root perimeter, convex area and solidity were observed with survivor zone root crowns. These results are consistent with earlier observations where a large number of lateral roots in survivor alfalfa plants were observed, however quantification of the root architectural changes was achieved with the current study (King 1923). This increase may be attributed to the emergence of an increased number of lateral roots following damage to the taproot and does not reflect greater depth that is associated with a tap root. Likewise, survivors had a greater number of root tips and holes than asymptomatic root crowns, which is another indication for root complexity. The observance of greater frequency of fine and medium roots in survivors may be due to the loss of apical dominance of the taproot resulting in the development of finer roots from the crown or damaged taproot, known as compensatory root growth (Rubio and Lynch 2007). Of note, more than half of the root crowns sampled from survivor zone lack taproots. On the contrary, root crowns from the asymptomatic zone exhibited significantly higher average and median root diameters suggesting that they possess proportionally thicker roots compared to the root crown from the survivor zone. Previous studies comparing greenhouse grown 18-28 days old alfalfa plants inoculated with *Pythium* spp. to non-infected plants showed a reduction in root system size and a lower degree of branching (Larkin et al. 1995). However, such studies focused on recently inoculated plants unlike the current study, which involved four-year old field grown alfalfa plants that have sustained PRR disease stress. Another plausibility for development of complex root system is the accessibility of more soil resources to the survivors owing to sparse stand in the survivor zone from loss of plants due to PRR. Similar changes in root features such as increased lateral root number and greater taproot diameter have been observed with greater spacing between alfalfa plants (Lamb et al. 2000). Although survivors had a complex root system compared to asymptomatic plants, this did not affect the computation time required to extract the suite of features from the digital images.

Roots by the nature of being in the ground, are inherently difficult to study without using destructive measures. We utilized the RhizoVision Crown phenotyping platform and showed that alfalfa plants surviving from PRR disease stress have root systems acclimated to damage caused by the disease. Despite the loss or damage to the taproot, survivors overcame this stress by developing additional crown and lateral roots contributing to a root system that was more complex than the asymptomatic roots, indicating an interesting interaction between soil microbes and plant architecture. Future research will need to address whether the acclimation of the diseased plant by increasing carbon allocation to crown and lateral roots is truly adaptive, and if so, whether the ability to acclimate by compensatory root growth could be a breeding target for increasing disease resistance. The methods used in this study could be applied to roots of crop plants undergoing other biotic, abiotic stresses or symbiotic interactions associated with the phytobiome.

The concept of low-input agriculture and the spatiotemporal dissimilarities in the distribution of water and nutrient sources in the soil have led in the identification of traits that have reduced metabolic cost for soil exploration (Lynch 2015; Lynch et al. 2014). The alfalfa plants from the survivor zone showing alternate root architecture under PRR disease stress presents a unique opportunity to further explore if this mode of survival has improved resource efficiency that would compensate for the loss of taproot and additional costs incurred by the plant while generating lateral and crown roots. Currently very limited options are available to manage PRR disease with no known alfalfa resistant varieties. Looking forward, it remains an open question whether the strategy of developing a different root architecture put forth by survivors is a heritable trait that could be used in resistance breeding and if these plants will be as persistent under other biotic and abiotic stress conditions.

## DATA AVAILABILITY

Root crown images and R statistical analysis code generated from this study are available on Zenodo.

York, Larry M., Young, Carolyn A., Mattupalli, Chakradhar, & Seethepalli, Anand. (2018). Images and statistical analysis of alfalfa root crowns from inside and outside disease rings caused by cotton root rot (Version 1.0.0) [Data set]. Zenodo. http://doi.org/10.5281/zenodo.2172832

## ACKNOWLEDGEMENTS

We thank the Noble Research Institute, LLC for funding this project.

